# Examination of genome-wide ortholog variation in clinical and environmental isolates of the fungal pathogen *Aspergillus fumigatus*

**DOI:** 10.1101/2022.03.23.485522

**Authors:** Maria Augusta Horta, Jacob Steenwyk, Matthew E. Mead, Luciano H. Braz dos Santos, Shu Zhao, John G. Gibbons, Marina Marcet-Houben, Toni Gabaldón, Antonis Rokas, Gustavo H. Goldman

## Abstract

*Aspergillus fumigatus* is both an environmental saprobe and an opportunistic human fungal pathogen. Knowledge of genomic variation across *A. fumigatus* isolates is essential for understanding the evolution of pathogenicity, virulence, and resistance to antifungal drugs. Here, we investigated 206 *A. fumigatus* isolates (133 clinical and 73 environmental isolates) aiming to identify genes with variable presence across isolates and test whether this variation was related to the clinical or environmental origin of isolates. The PanCore genome of *A. fumigatus* constitutes 13,085 ortholog groups, of which 7,773 (59.4%) are shared by all isolates (CORE) and 5,312 (40.6%) vary in their gene presence across isolates (ACCESSORY). Despite differences in the distribution of orthologs across all isolates, no significant differences were observed among clinical vs. environmental isolates when accounting for phylogeny. Orthologs that differ in their distribution across isolates tend to occur in low frequency and/or be restricted to specific isolates; thus, the degree of genomic conservation between orthologs of *A. fumigatus* is high. These results suggest that differences in the distribution of orthologs within *A. fumigatus* cannot be associated with the clinical or environmental origin of isolates.

**Importance:** *Aspergillus fumigatus* is a cosmopolitan species of fungi responsible for thousands of cases of invasive disease. Clinical and environmental isolates of *A. fumigatus* exhibit extensive phenotypic differences, including differences related to virulence and antifungal drug resistance. A comprehensive survey of the genomic diversity present in *A. fumigatus* and its relationship to the clinical or environmental origin of isolates can contribute to the prediction of the mechanisms of evolution and infection of the species. Our results suggest that there is no significant variation in ortholog distribution between clinical and environmental isolates when accounting for evolutionary history. The work supports the hypothesis that environmental and clinical isolates of *A. fumigatus* do not differ in their gene contents.

## Introduction

*Aspergillus fumigatus* is a soil-borne and ubiquitously distributed filamentous fungus that recycles organic matter. *A. fumigatus* is also a major opportunistic human pathogen that causes life-threatening invasive pulmonary aspergillosis (IPA) in immunocompromised hosts. *A. fumigatus* is responsible for an estimated 3,000,000 cases of aspergillosis annually and more than 200,000 cases of IPA each year, reaching a mortality rate of up to 90% in the most susceptible populations (Brown et al., 2012; Bongomin et al., 2017; Firacative, 2020). One reason for the prevalence of *A. fumigatus* as the main clinical etiological agent of aspergillosis is its cosmopolitan distribution and high prevalence in the environment, with the average human inhaling hundreds of airborne asexual spores (conidia) of *A. fumigatus* daily (Latgé &Chamilos, 2020; Wéry, 2014).

Genetic heterogeneity of *A. fumigatus* has been extensively described (Debeaupuis et al., 1997; Frías-De León, 2011; Kowalski et al., 2016; Lind et al., 2017; Zhao & Gibbons, 2018; Barber et al; 2021; Kowalski et al., 2021; Lofgren et al., 2021). *A. fumigatus* also exhibits phenotypic heterogeneity in virulence in animal models of IPA (Kowalski et al., 2016; Ries et al., 2019) and in its drug resistance profiles to antifungals (Colabardini et al., 2021). Azoles are the first-line treatment against IPA, but the recent emergence of azole-resistance in *A. fumigatus* has made treatment even more challenging (Latgé &Chamilos, 2020). Resistance to azoles is an important concern and global surveillance studies reveal that 3.2% of *A. fumigatus* isolates are resistant to one or more azoles, with large discrepancies between geographic regions (van der Linden et al., 2015). The azole target is the 14-α-sterol demethylase encoded by *cyp51A*, and the main mechanisms of resistance found in environmental and clinical isolates involve mutations in the *cyp51A* coding region and tandem duplications in its promoter region (Dudakova et al., 2017; Latgé &Chamilos, 2020). The genetic heterogeneity of the *A. fumigatus* population to azole resistance has been described (Dudakova et al., 2017; Barber et al., 2020b; Lofgren et al., 2021).

Several recent studies explored the genetic diversity across *A. fumigatus* isolates, with some of these studies aiming to explain how the observed genomic diversity contributes to variation in secondary metabolism, virulence, and antifungal drug resistance (Lind et al., 2017; Zhao & Gibbons, 2018; McCarthy &Fitzpatrick, 2019; Barber et al., 2021; Lofgren et al., 2021). Zhao and Gibbons (2018) looked at gene copy number variation across *A. fumigatus* isolates and found gene presence-absence polymorphisms associated with virulence. Lofgren et al. (2021) analyzed gene sequence and gene presence-absence variation to support the existence of three distinct *A. fumigatus* populations; specific alleles and genes involved in drug resistance were often distributed in a population-specific manner. Barber and collaborators (2021) described the genomic variation associated with clinical isolates related to triazole resistance and virulence factors. Isolates were assigned to genetic clusters based on their SNP profiles and placed in the phylogeny, thus determining clusters enriched in clinical isolates, as well the functional genomic profiles of the isolates from the clusters (Barber et al., 2021). Another recent study determined the population structure of azole-resistant isolates from environmental and clinical sources observing signatures of positive selection in both regions containing canonical genes encoding fungicide resistance in the ergosterol biosynthetic pathway as well as in regions that have no defined function (Rhodes et al.,2021).

Understanding the genetic divergence across isolates is essential to study the genetic mechanisms underlying specific phenotypes, including differences in virulence and pathogenicity across *A. fumigatus* isolates. Here, we investigated if variation in ortholog gene content in a population of *A. fumigatus* isolates is associated with isolate origin (clinical or environmental). To do so, we inferred orthologous groups of genes encoded in the genomes of 206 globally distributed clinical and environmental isolates of *A. fumigatus*. Despite notable differences in orthogroup gene content across all isolates, no significant correlation between gene distribution and the origin of isolates was detected, pointing to similar orthogroup content between all clinical and environmental isolates when accounting for phylogeny. These results support the hypothesis that clinical and environmental isolates of *A. fumigatus* do not differ in their gene content.

## Results

### Orthology inferences among isolates

The examination of orthologous gene relationships among genes of all *A. fumigatus* genomes (genome metrics are presented in Supplementary Material 1) revealed a PanCore genome composed of CORE groups (i.e., genes present in all isolates) and ACCESSORY (AC) groups (i.e., accessory genes present in one or more, but not all, isolates). It represents 13,085 orthogroups, those 7,773 (59.4% of the PanCore genome, corresponding to 1,606,893 genes) are CORE groups and 5,312 (40.6% corresponding to 251,058 genes) are AC groups (Supplementary Material 2). The number of orthogroups and predicted sequences classified for each isolate was also investigated to reveal the variation across isolates genomes and their origin (Figure 1). Thus, we determined the frequency of isolates that share AC groups: 860 AC groups were identified in >95% of the isolates (6.6% of the PanCore genome), 774 groups in 5–95% of the isolates (5.9% of the PanCore genome), and 1,021 AC groups are present in fewer than 5% of the isolates (7.8% of the PanCore genome), showing the low variation in the distribution of orthologs across genomes (Supplementary Material 2). In addition, 2,675 genes were isolate-specific, hereafter referred to as singletons, which represent 0.14% of the total number of genes of the dataset and are exclusively found in 123 isolates (Supplementary Material 3).

**Figure 1:**
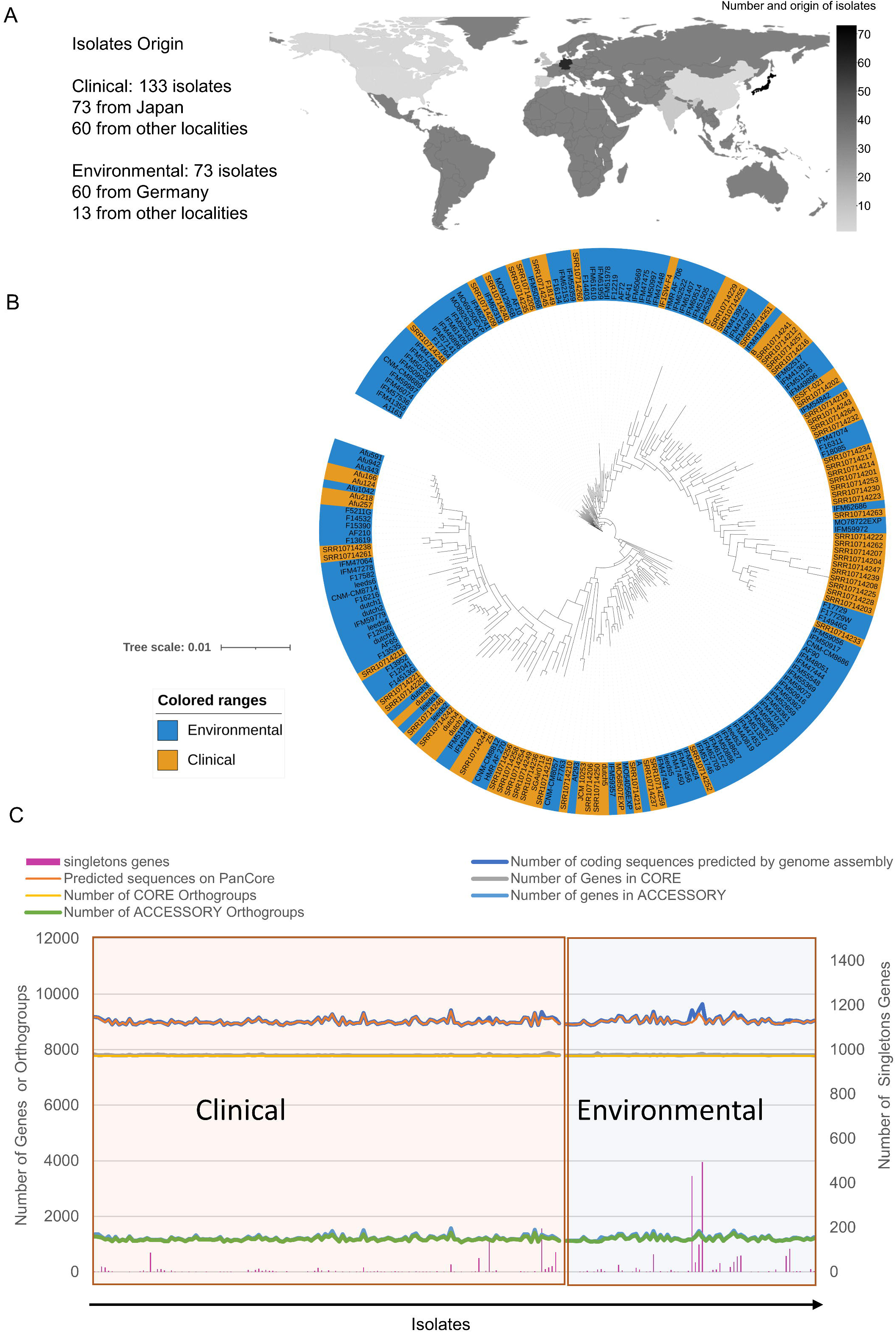
Isolates comparison on PanCore genome analysis, showing the high similarity of genomes in terms of phylogenetic distribution and gene classification, despite the worldwide distribution of isolates. A) Geographical distribution of the isolates investigated showing the country origin, the color dimming of countries represents an increase in the number of isolates. B) Phylogenomic analysis of *A. fumigatus* isolates reveals clinical and environmental isolates do not belong to distinct clades. C) Total numbers determined per isolate of clinical and environmental origin, X-axis represents isolates and right Y-axis line plot represents the number of genes (genes classified on CORE and AC orthogroups per isolate, as well the number of predicted genes for each isolate) or orthogroups (number of CORE and AC orthogroups present at each isolate); left Y-axis bar plot represent the number of singletons genes determined in each isolate.

### Functional classification of PanCore genome

The total number of genes in the PanCore genome was examined (Supplementary Material 1) and compared with Af293 and A1163, the most well annotated and commonly used reference isolates for *A. fumigatus* (Nierman et al., 2005; Fedorova et al., 2008). The average number of predicted genes in each genome was 9,023 ± 118 genes; the genomes of clinical and environmental isolates encoded 9,008 ± 101 and 9,051 ± 140 genes, respectively. The frequency of CORE and AC groups was similar across isolates, with a slightly higher average number of genes for the environmental set of isolates (CORE: environmental 9,027 ± 101, clinical 9,001 ± 93; AC: environmental 1,223 ± 95, clinical 1,205 ± 88). Notably, the average number of genes in the entire dataset was nearly the same as the number of genes in the commonly used reference strains Af293 and A1163 (Supplementary Table 1). For the total dataset, the average number of CORE genes was 9,010 ± 97 and of AC genes was 1,211 ± 91 genes. To identify variation in genome content between isolates, the orthogroups were further classified using eggNOG v5.0 (Huerta-Cepas et al., 2019) and confirmed by the function of orthogroup members from clinical isolates Af293 and A1163 (Nierman et al., 2005; Fedorova et al., 2008) (Supplementary Material 4). For Af293, a total of 86.3% of the genes in its genome were CORE genes and 13.7% were AC genes.

Comparison of the ontology terms among genes present in CORE and AC orthogroups revealed differences in functional classifications (Supplementary Figure 1). The “molecular adaptor activity” (GO:0060090) was present in the CORE group but absent in the AC group. The protein classification with the largest proportion in the CORE and AC group was “metabolite interconversion enzyme” (PC00262), and the protein class with the largest absence in AC group is “nucleic acid metabolism protein” (PC00171). In the AC group, there is a smaller proportion of peptidases and hydrolases acting on acid anhydrides (GO:0008233 and GO:0016817, respectively) than the CORE group; in contrast, in the AC group, there is a larger proportion of hydrolases acting on glycosyl bonds and ester bonds (GO:0016798 and GO:0016788, respectively). Notably, higher relative proportions of oxidoreductase activity orthogroups were observed among the AC groups—specifically for “dioxygenase activity” (GO:0051213) and “oxidoreductase activity acting on peroxide as acceptor” (GO:0016684). These results demonstrate that CORE and AC groups of genes functionally differ.

The annotation of the Af293 genome facilitated prediction of protein products encoded by 8,631 (66.0%) orthogroups (Supplementary Material 4). Moreover, orthogroups putatively associated with *A. fumigatus* adaptability, survival, and virulence—which are hereafter referred to as “Target Genes” (Figure 2 and Supplementary Material 5)—could easily be identified. Target genes include kinases, conidiation-related genes, G-protein coupled receptors (GPCRs), ergosterol biosynthesis enzymes, phosphatases, ABC (ATP-binding cassette) transporters, major Facilitator Superfamily (MFS), transporters, secondary metabolites (BGCs), transcription factors, and genetic determinants of virulence. Across 1,358 orthogroups, we identified a total of 1,566 target genes, which spanned 188 AC and 1,170 CORE groups (Figure 2A).

**Figure 2:**
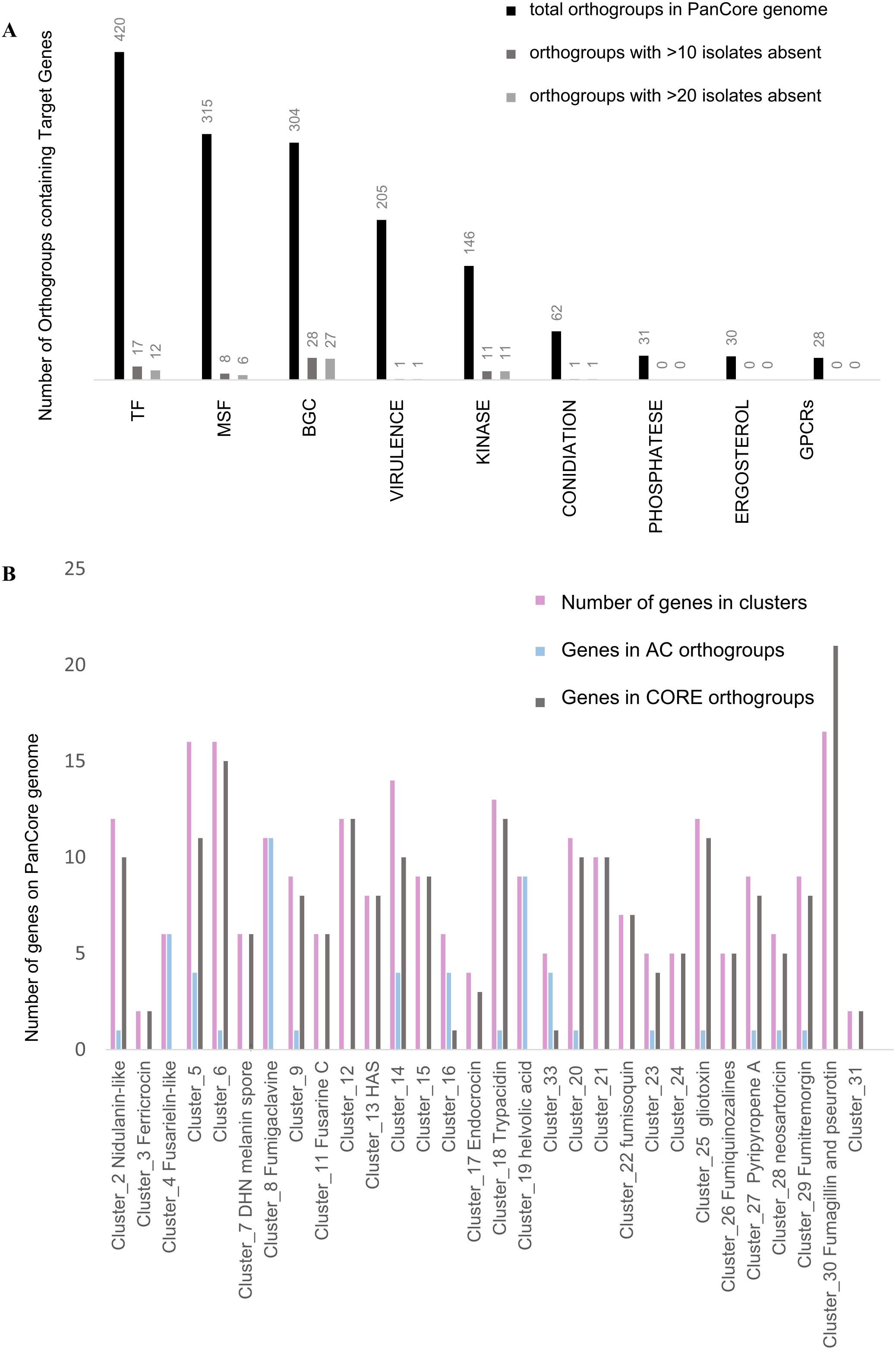
Analysis of Target Genes across *A. fumigatus* isolates shows that the variation on orthologous genes distribution across isolates is a low frequency event that usually affects less than 10 isolates (>5%), maintaining the majority of Target Genes classified on CORE orthogroups. A) Genes putatively associated with *A. fumigatus* adaptability, survival, and virulence (or Target Genes) were localized in PanCore orthogroups and classified according their function. B) Gene presence and absence patterns among genes encoded in BGCs. TF: transcription factor; MFS: Major facilitator superfamily transporter; BGCs: Biosynthetic Gene Clusters.

GPCRs, ergosterol biosynthesis genes, conidiation genes, and phosphatases were observed among CORE groups. In contrast, genes associated BGCs, kinases, TFs, and MFSs were observed among AC groups. For example, PkaC/Afu2g12200 (the catalytic subunit of cAMP-dependent protein kinase) is present in a single-copy across all isolates whereas Fhk2/Afu3g07130 (a two-component response regulator) is present in 101 isolates in single-copy and present in multiple copies in 105 isolates (Supplementary Material 5).

To determine variation among genes related to secondary metabolite production, we considered the biosynthetic gene cluster (BGCs) classification from Lind *et al*. (2017) that described the Af293 genetic clusters and the predicted functions for a partial number of clusters. For our analysis, we considered only those clusters where at least 80% of predicted genes were present as a biosynthetic gene cluster (BGC) in each isolate respecting the distribution of BGCs as previously reported (Lind et al., 2017). According to this criterion, BGCs 1, 10 and 33, which biosynthesize unknown secondary metabolites, are not conserved and were excluded (Figure 2B, Supplementary Material 5). A total of 270 genes from BGCs were identified on the orthogroups, of which 81.4% are in CORE and 18.5% are in AC groups. The gliotoxin BGC is conserved and 12 of 13 genes are CORE genes, and only isolate IFM59779 lacks the orthologous genes to *gliK* (Afu6g09700). To further support this finding, we mapped the IFM59779 reads onto the Af293 genome and found that reads did not map to *gliK* supportive of gene loss (Supplementary Figure 2). The same conservation patterns were observed for helvolic acid (Cluster 19) and nine genes were identified among CORE and AC groups, respectively. Of note, the helvolic BGC was absent in CNM-CM8812 suggestive of an isolate-specific loss of genomic content.

### Gene copy number variation

Another important aspect that may contribute to organismal adaptive evolution and the evolution of fungal pathogens is the variation on gene copy number (Yang et al., 2012; Steenwyk et al., 2016; Fernández &Gabaldón, 2020). To evaluate gene copy number variation, we examined the number of genes assigned for each isolate per orthogroup. Multi-copy genes were observed at a low frequency across isolates. Specifically, in 8.4% of the PanCore genome (1,094 groups, 478 AC and 616 CORE) there was at least one isolate with multiple gene copies and in 1.1% (146 orthogroups, 78 AC and 68 CORE) of the PanCore genome 5% of isolates (i.e., more than 10 isolates) had two gene copies. Orthogroups with isolates that encoded three gene copies were observed in 1.5% of the PanCore genome (194 groups, 133 AC and 61 CORE). In only 14 groups (0.1% of the PanCore genome, 13 AC and 1 CORE), three gene copies were observed in 5-23% of isolates (Supplementary Material 2).

We next evaluated gene copy number variation among orthogroups presenting multi-copy genes in more than 5% of isolates (Figure 3A, Supplementary Material 2). In addition, we detailed the variation of two copies genes according isolates origin for the Target Genes, genes putatively associated with *A. fumigatus* adaptability, survival, and virulence (Figure 3B, Supplementary Material 5). Genes present in two or three copies were observed among 14 orthogroups with varying frequency across isolates—more specifically, 9.7 to 42% of isolates had two genes present in an orthogroup and 5.3 to 23% had three genes present in an orthogroup. Among these 14 orthogroups, OG0000013, which encodes a putative amino oxidase, is present in the CORE group. The remaining 13 are observed in the AC group and encode genes with diverse functional categories and domains: predicted FAD-binding domain, transposition, RNA-mediated and Chromo (CHRromatin Organisation MOdifier) domain protein function, no protein function classification (unknown function DUF3435, conserved hypothetical protein) or predicted FAD-binding domain, transposition, RNA-mediated and Chromo (CHRromatin Organisation MOdifier) domain protein function. Notably, the orthogroup OG0000009, which was observed in the AC group, has a putative protein function related to the constitution of regulatory subunit 26S proteasome, the major non-lysosomal protease in eukaryotic cells. This multiprotein complex is involved in the ATP-dependent degradation of ubiquitinated proteins, playing a key role in the maintenance of protein homeostasis by removing misfolded or damaged proteins (Bard et al., 2018). Among all isolates, 49% two gene copies (39% environmental and 61% clinical isolates) and 5% had three copies (89% environmental and 11% clinical) of OG0000009. Two gene copies were also observed in orthogroups encoding TF genes—for example, the 27% of isolates had two copies assigned to orthogroups OG0000044 (Forkhead transcription factor Fkh1 2) and OG0000047 (C6 finger domain protein). Of note, the distribution of isolates by origin in both orthogroups is different from the distribution across the whole dataset (35.5% environmental and 64.5% clinical isolates across all orthogroups). Specifically, in OG0000044 and OG0000047 two gene copies were present in 82% and 87.5%, respectively, among environmental isolates whereas two gene copies were observed among 18% and 12.5% of clinical isolates (Supplementary Material 2).

**Figure 3:**
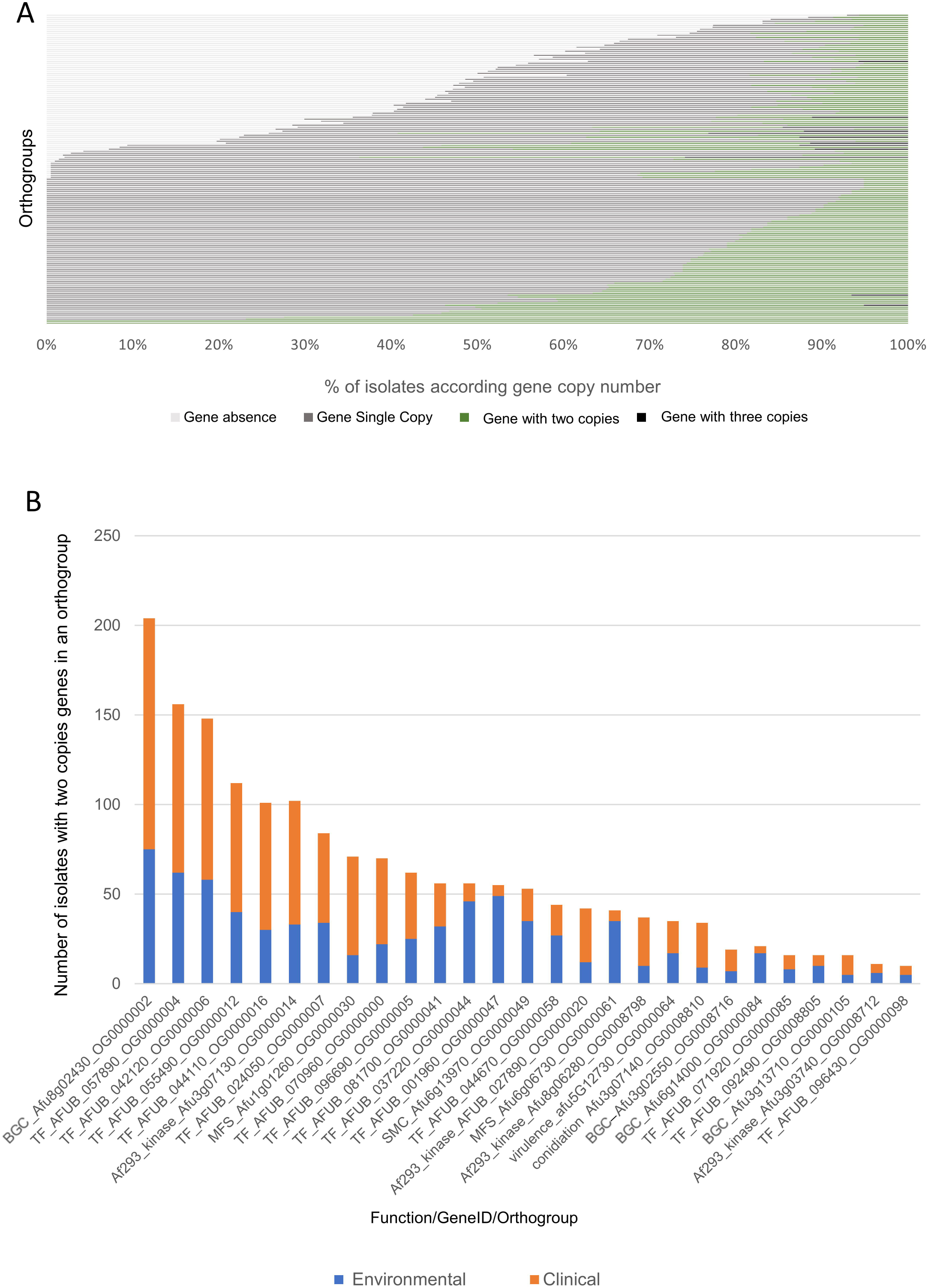
The copy number variation detected in PanCore genome shows differences in the proportion of absent and multiple copies genes across orthogroups, and on the distribution of the isolates within two copies genes. A) Distribution of isolate presence and absence per orthogroup. Genes present in single copy are shown in grey, genes present in two copies are shown in green, genes present in three copies are dark grey, genes that are absent are light grey. B) Target genes that presents two copies of genes and their differences on the distribution of isolates within orthogroups according to clinical or environmental origin.

Zinc finger transcription factors often control the expression of multiple genes and trigger control cascades that dramatically influence the regulation of abiotic stresses (Han et al., 2020), including influencing cation homeostasis and the detoxification process in *A. nidulans* (Spielvogel et al., 2008). Two orthogroups that encode zinc finger TFs had high frequency of isolates presenting two copies of the genes —specifically, 72.8% and 50.5% of isolates, respectively on orthogroups OG0000006 and OG0000016 (Supplementary Material 2). However, the frequency of isolates that encoded two gene copies according to the origin was similar to the background. Specifically, 37.0% and 63.0% of environmental and clinical isolates, respectively, had two gene copies present in orthogroup OG0000006; 29.0% and 71.0% of environmental and clinical isolates, respectively, had two gene copies present in orthogroup OG0000016.

Another gene often present in multiple copies are signal sensor histidine kinases. For orthogroups OG0000012 and OG0000014, 55.8% and 49% of isolates had two gene copies in each orthogroup, respectively (Supplementary Material 2). In the opportunistic human yeast pathogen *C. albicans*, these proteins encode two-component signaling proteins that transmits signals through the high-osmolarity glycerol response 1 (HOG1) and mitogen-activated protein kinase (MAPK cascade) in response to environmental osmotic stimuli (Brewster et al., 1993, Kruppa &Calderone, 2006). In *A. fumigatus*, high-osmolarity glycerol (HOG) and cell wall integrity pathways are important for the adaptation to different forms of environmental adversity such as osmotic and oxidative stresses, nutrient limitations, high temperatures, and other chemical and mechanical stimuli (Silva et al., 2020, Assis et al., 2020, Manfiolli et al., 2019). The frequency of two gene copies with respect to isolate origin was similar to the frequencies of environmental (35.5%) and clinical (64.5%) isolates across the whole dataset, suggesting no differentiation of multiple copies according to origin. Overall, the majority of the orthogroups (68.7%, 8,997 orthogroups) of the PanCore genome had a similar distribution of isolate origin of the dataset (64.5% clinical and 35.5% environmental), considering a standard deviation of 5% on isolates distribution.

We next tested whether increased copy number was observed among Target Genes, in which multiple copy genes detected in more than 5% of isolates are in orthogroups that encode TFs, BGCs and kinases respectively 12, 8 and 5 orthogroups (Figure 3B, Supplementary Material 5).

To determine if gene copy number per orthogroup varied between environmental and clinical isolates, we conducted a phylogenetic ANOVA test (phylANOVA) (Garland et al., 1993). To do so, single copy orthologs were first used to infer the evolutionary history of the 206 isolates (Figure1A and 1B, unrooted tree provided in Supplementary Figure 3). In these, we did not observe a significant number of associations between the phylogenetic position of an isolate and its status of clinical or environmental origin (p-value > 0.05; phylANOVA). These findings suggest that when accounting for evolutionary history, environmental and clinical isolates do not substantially differ in gene copy number per orthogroup. Corroborating with this observation, the number of classified sequences on PanCore genome and the orthogroups classified for each isolate maintains a linear trend across clinical and environmental isolates (Figure 1C).

### Origin of singleton

To investigate the possible origin of the orthogroups of singletons genes, each sequence was BLASTed against fungal and bacterial genes from the nonredundant protein database (nr-NCBI). The “Sugar transporter (major facilitator superfamily (TC 2.A.1.1))” and “Fungal specific TF domain” werethe functional descriptions most frequently observed best BLAST hit. Mainly the singletons genes contain sequences with orthologs in fungi and sequences with a high degree of differentiation, with orthologs in a different genus or no ortholog at all (Supplementary Material 3).

Taken together, these findings highlight the conservation of the distribution of orthologous genes, with a gene frequency occurrence 40.5% for AC genes distributed across *A. fumigatus* dataset.

### Transposable elements

Mobilization of transposons can cause a variety of localized effects on genomes such as gene inactivation or modification, and they are often found in multiple, highly conserved copies. Transposons also represent potential ‘hotspots’ for the occurrence of inversions, translocations, deletions and gene copies (Muszewska et al., 2019). Two non-functional type I transposons have been reported in *A. fumigatus*, AfutI and AfutII (Neuveglise et al., 1996, Paris &Latgé, 2001), a seemingly non-functional class II element Taf1 (Monroy &Sheppard, 2005) and Aft1, a type II transposon of the Tc1/Mariner superfamily (Hey et al., 2007). The search for transposons Aft1 and Taf1 in all *A. fumigatus* genomes was performed considering nucleotide identity (Supplementary Material 6). This analysis revealed variation in the number of repeated elements over the genomes. It has been reported that Taf1 is present in different locations and copy number among clinical isolates of *A. fumigatus* (Monroy &Sheppard, 2005; Hey et al., 2007). We identified 20 copies of Aft1 (EF100757.1, 1882 bp) and 13 copies of Taf1 (AY971670.1, 2,023 bp sequence) with nucleotide identity >99.0% in the isolate Af293 (Supplementary Material 6). Aft1 was found in 190 isolates in 1 to 20 copies and Taf1 was present in 204 isolates varying from 1 to 18 copies (Supplementary Figure 4, Supplementary Material 6). The isolate with the highest number of copies for both elements was the environmental strain SGAir0713 (18 copies of Taf1; 21 copies of Aft1). TAf1 orthologs were not identified in the environmental isolates B-1-26-5 and C-1-27s-1. Following findings from previous studies (Muszewska et al., 2019) (Schrader &Schmitz, 2019), we hypothesize that transposable element variation may be associated with phenotypic variation following previous.

## Discussion

*Aspergillus* taxonomic sections show extensive variation and the traits and genetic elements that contributed to this variation are still under investigation (Rokas &Galagan, 2008; Dos Santos et al., 2020; Rokas et al., 2020). Previous comparative genomic investigations in the *Aspergillus* genus characterize both species diversification and variation within species (Vesth et al., 2018). The first study that investigated the distribution of orthologous genes in *A. fumigatus* used 12 isolates and identified 8,073 CORE groups and 1,964 AC groups (McCarthy &Fitzpatrick, 2019). The increasing availability of sequencing data has allowed for even more precise description of the *A. fumigatus* PanCore genome (Lofgren et al., 2021; Barber et al., 2021). The largest PanCore genome analysis among 300 *A. fumigatus* isolates (83 clinical and 217 environmental, the majority from Germany) revealed the *A. fumigatus* PanGenome is much larger than previous estimates (Barber et al., 2021). Previous analyses (Barber et al., 2021) suggested the PanGenome is closed. Corroborating these findings, we inferred a closed PanGenome and identified comparable numbers of total orthogroups (including CORE and AC orthogroups), which may be in part due to partial overlap between isolates used in both studies; however, we note our dataset contained approximately twice the number of clinical isolates. Both studies report infrequent variation, fewer than 5% of the isolates among genes related to virulence (Figure 2; Supplementary Material 2) (Barber et al., 2021). Nevertheless, the distribution of orthologs cannot be directly compared between the studies because Barber and collaborators did not identify individual genes on orthogroups. Thus, the distribution of orthologous genes provided here contains important information to the *Aspergillus* research community (Supplementary Material 2), as well the possibility to identify Af293 orthologs on the PanCore genome (Supplementary Material 4).

Across all isolates, we found that *A. fumigatus* harbors variation in terms of the number of total predicted protein coding genes ranging between 8,857 and 9,638 (for the clinical isolate IFM59779 and environmental isolate C-1-33L-1, respectively) (Supplementary table 1). The isolates with more genes classified in the PanCore genome were the clinical isolate MO78722EXP and the isolate ISSFT-021, which was obtained from the international space station, with 9,117 and 8,972 genes classified respectively. In the AC set, the clinical isolates IFM59361 and 12-7504462 had the highest number of AC classified genes, 1,570 and 1,521, respectively (Supplementary Material 7), revealing variation on gene copy numbers (absence, single or multiple gene copies) across isolates (Figure 3A). Other studies suggest that the genetic variants— SNPs, indels, and gene presence absence polymorphisms—across *A. fumigatus* isolates may provide evidence of distinct populations of *A. fumigatus* (Lofgren et al., 2021).

We investigated the distribution of genes with functional associations with relevant pathways investigated in *A. fumigatus* studies in the PanCore genome, focusing on important mechanisms significant to the phenotypic differentiation of isolates (Supplementary Material 5). Our analysis revealed extensive variation in copy number among genes encoding GPCRs, phosphatases, ABC-transporters, kinases, TFs, MSF-transporters, and proteins important for conidiation, virulence, and secondary metabolite production (Figure 2A). Notwithstanding these differences, gene copy number per orthogroup did not differ between clinical and environmental isolates.

*A. fumigatus* produces a diversity of secondary metabolites (SM) and efflux pumps that serve as defense systems (Keller et al., 2005) (Kwon-Chung &Sugui, 2013). In fungi, the genes in pathways that synthesize SMs are typically located next to each other in the genome and organized in contiguous gene clusters (BGCs) (Clevenger et al., 2017; Macheleidt et al., 2016, Lind et al., 2017; Rokas et al., 2018). The gliotoxin BGC impacts *A. fumigatus* virulence and is widely produced by *Aspergillus* species (Steenwyk et al., 2020; Mead et al.,2021). Here, we observed the conservation of the gliotoxin BGC across the PanCore genome (Figure 2B). However, other BGCs, such as fumitremorgin, presented heterogeneity on the genetic arrangement of BGCs within species (Steenwyk et al., 2020), an observation that is typified by the total absence of these genes in the environmental isolate B-1-70s-1 (Supplementary Material 5). From the genes belonging to BGCs, 27 presented significant variation in species distribution in the PanCore genome (Figure 2B). All genes from the helvolic acid BGC were absent from isolate CNM-CM8812. All genes in BGC 4, predicted to produce a Fusarielin-like metabolite, were classified as accessories and are absent in different isolates, such as A1163 (ASM15014v1) that lost all genes from this BGC. The fumitremorgin BGC, Cluster 29, presented a unique isolate (Afu343) with two copies of all genes of the BGC. Of note, our analysis examined the presence and absence of genes encoded in BGCs, but further examination of physical clustering is warranted. Nonetheless, our findings corroborated previous descriptions of variation among BGCs within this species (Zhao & Gibbons, 2018; Lind et al., 2017; Barber et al., 2021).

The genetic diversity across species in virulence and drug resistance mechanisms have been extensively reviewed (Kwon-Chung &Sugui, 2013; Rokas et al., 2020; dos Santos et al., 2021). Among clinical isolates of *Aspergillus* species, species- and isolate-specific polymorphisms were reported in the 14α-sterol demethylase gene *cyp51A* (Afu4g06890) and in the 1,3-beta-glucan synthase catalytic subunit gene *fks1*, target genes for azoles and echinocandins, respectively (Dudakova et al., 2017; Zhao & Gibbons, 2018; Dos Santos et al., 2020). We observed variation in gene copy number and sequence for *cyp51A* (Afu4g06890/orthogroup OG0003434), *cyp51B (*Afu7g03740/ orthogroup OG0000331) and *fks1* (Afu6G12400/ orthogroup OG0003140), and these were all members of CORE groups (Supplementary Material 2/ Supplementary Material 4).

The genetic determinants of *A. fumigatus* virulence were previously described (Mead et al., 2021) and appear as a conserved set of genes on our PanCore genome. From 204 studied genes, only 23 were classified in AC groups. The *pes3* gene, encoding a nonribosomal peptide synthetase 8 (Afu5G12730) is localized in the CORE orthogroup OG0000064 and presents two copies for 38 isolates (44% environmental and 55% clinical). The AC group OG0008770 contains the important factor for virulence *cgrA* (Afu8G02750/Nucleolar rRNA processor) and shows strong variation: 50 isolates do not contain the gene (82% environmental and 18% clinical) and 156 isolates have a single copy of the gene (21% environmental and 79% clinical), suggesting a tendency to lose this gene in environmental isolates. The variation in AC is mainly due to the absence of orthologs in one or two isolates, and CNM-CM8812 lacked orthologous genes in nine orthogroups encoding genetic determinants of virulence.

Conidiation is critical for *A. fumigatus* dispersal to new environments, including the human lung (Stewart et al., 2020). FluG (Afu3g07140; OG0008810) acts upstream of the BrlA activator of conidiation (Ni et al., 2010; Adams et al., 1988) and was absent in 99 isolates (41% environmental and 54% clinical), present in single copy in 79 isolates (32% environmental and 64% clinical), and present in two copies in 35 isolates (24% environmental and 73% clinical). The extensive absence of *FluG* orthologs suggests that FluG may not be essential for conidiation.

In environmental isolates, the impact of fungicides on the number of gene duplications in *A. fumigatus* is variable and the overall resistance frequency among agricultural isolates is low (Barber et al., 2020b; Latgé &Chamilos, 2020) relative to the increasing azole resistance trend in clinical *A. fumigatus* isolates (Buil et al., 2019). Isolates collected from soil after the growing season and azole exposure show a subtle but consistent decrease in susceptibility to medical and agricultural azoles (Barber et al., 2020b). Whole-genome sequencing indicates that despite variation in antifungal susceptibility, fungicide application does not significantly affect the population structure and genetic diversity of *A. fumigatus* in the fields (Barber et al., 2020a), consistent with our finding of conservation in orthogroup distribution among clinical and environmental isolates.

PanCore genome analysis allowed us to detect instances of copy number variation, especially when multiple gene copies were present. For example, among 56 isolates that encode two copies of the TF encoding genes Afu1g01560 (OG0000047) and Afu3g11960 (OG0000044), 87.5% and 82%, respectively, are environmental isolates whereas 12.5 % and 18%, respectively, are clinical isolates in two copies. Two genes encoding kinase activity— Afu8g06280 (OG0008798) and Afu3g07130 (OG0000014)—tended to be more widely observed among clinical isolates, 73.6% and 69% were clinical isolates in two copies, whereas 26.3% and 31% are environmental isolates, respectively. The observed variation qualitatively differed from the variation observed across all orthogroups (35.5% environmental and 64.5% clinical). We also highlight transposable elements *Taf1* and *Aft1* as a source of substantial genetic variation among *A. fumigatus* isolates.

Our work investigated variation in orthogroup content among a comprehensive panel of clinical and environmental isolates of *A. fumigatus*. Despite identifying substantial variation in gene presence, absence, and copy number across the whole dataset, no variation was observed among environmental and clinical isolates when accounting for phylogeny. This finding raises the hypothesis that environmental and clinical isolates may differ due to other types of genetic variation (e.g., SNPs and indels) or do not substantially differ at all.

## Materials and Methods

### Dataset assembly

The dataset was constructed based on genomes previously referenced (Zhao et al., 2020; Steenwyk et al., 2020; Barber et al., 2020b). The isolate descriptions are provided in Supplementary Material 7. Three principal datasets were used (details are in Supplementary Material 1): 1) Clinical isolates from different Japanese cities were provided through the National Bio-Resource Project (NBRP), Japan (http://nbrp.jp/) (NCBI BioProject PRJNA638646), and additional isolates (Takahashi-Nakaguchi et al., 2015) from NCBI BioProject PRJDB1541 originated from different patients, sources and infections, described by (Zhao et al., 2020) (Zhao et al., 2019); 2) Public genomes available from clinical and environmental isolates were used, including Af293 (GenBank accession: GCA_000002655.1), CEA10 (strain synonym: CBS 144.89/ FGSC A1163; GenBank accession: GCA_000150145.1), HMR AF 270 GenBank accession: GCA_002234955.1), Z5 (GenBank accession: GCA_001029325.1), and additional isolates from Spain (Dos Santos et al., 2020), Portugal and other origins were assembled (Steenwyk et al., 2020; Steenwyk et al., 2021c) and public under NCBI BioProject PRJNA577646; 3) Environmental soil samples from German fields before and after fungicide application were used to isolate *A. fumigatus* isolates. The DNA extraction, genome sequencing and quality assessment is described by (Barber et al., 2020a), and the reads under BioProject number PRJNA595552 were trimmed using Trimmomatic, v0.36 (Bolger et al. 2014) and used as input to the SPAdes, v3.11.1 (Bankevich et al., 2012) to obtain the genome assemblies. The assemblies from studies 2 and 3 were kindly provided by the groups of prof. Rokas and Gibbons. The samples originated from 133 clinical and 73 environmental isolates. Clinical isolates are largely from Japan (73 isolates) and Europe (Zhao et al., 2020), and environmental isolates are from German (60 isolates) soil samples (Barber et al., 2020b) and other localities (Supplementary Material 7).

### Data Analysis

All genome assemblies were tested by quality metrics and scaffold size evaluation through QUAST (Gurevich et al., 2013) (Supplementary Material 1). A new *A. fumigatus* training model was constructed and used during the consensus prediction sequences process by the software AUGUSTUS, v3.3.2 (Stanke &Waack, 2003) for all assemblies. To determine gene-content completeness, we used the BUSCO v2.0.1 pipeline (Waterhouse et al., 2018) to examine assembly contigs for the presence of near-universally single-copy orthologs (Supplementary Figure 5). The *eurotiales* OrthoDBv10 dataset (Kriventseva et al., 2019) of near-universally single-copy orthologs was used. Our analyses identified genomes with a maximum of 20 missing BUSCO genes. These metrics suggest that all 206 *A. fumigatus* genomes are non-fragmented and suitable for comparative genomic analyses. Genome metrics are presented in Supplementary Material 1 and BUSCO classification for isolates genomes is presented at Supplementary Figure 5.

To identify gene families across genomes, we used a Markov clustering approach. Specifically, we used OrthoFinder, v2.3.8 (Emms &Kelly, 2019), with inflation parameter of 1.5. After normalizing BLAST bit scores, genes are clustered into discrete orthogroups using a Markov clustering approach (van Dongen 2000). The eggNOG-mapper was used to assign GO, KEGG and PFAM terms to orthogroups based on their orthology relationships (Huerta-Cepas et al., 2017). For that purpose, we selected the eukaryotic eggNOG database (euNOG). Protein-coding genes were further classified by Gene Ontology terms using Panther v16 (Mi et al.,2021).

### Matrix construction and Statistical Analysis

Based on OrthoFinder results, two matrices were constructed: 1) A matrix that described the genes in the orthogroups; 2) A matrix that describes the number of genes from each isolate in the orthogroups. The gene counting matrix established the number of absent, single copy or multiple copy genes per isolate (Supplementary Material 2). Based on the gene identification matrix, the gene IDs of target orthogroups were selected to obtain the respective coding sequences. The set of coding sequences were further annotated using eggNog5.0 (Huerta-Cepas et al., 2019) and the functional profile was constructed based on the annotation of the sequence. The statistical computing and data analysis was done using R (R Team, 2013). We considered orthogroups with variation in more than 5% of isolates to avoid orthogroups with false positive cases due to assembly issues.

To reconstruct the evolutionary history of all *A. fumigatus* isolates, we implemented a workflow similar to previously established protocols (Steenwyk et al., 2019). More specifically, we first individually aligned amino acid sequences of single-copy orthologs using MAFFT, v7.402 (Katoh &Standley, 2013), with the “auto” mode. Next, codon-based alignments were generated by threading the corresponding nucleotide sequences onto the amino acid alignments using the “thread_dna” function in PhyKIT, v1.2.1 (Steenwyk et al., 2021). The resulting codon-based alignments were trimmed using ClipKIT, v1.1.5 (Steenwyk et al., 2020b) with the “smart-gap” mode. To save computational time during strain tree inference, we subsampled all single copy orthologs for the 2,500 most phylogenetically informative according to the number of parsimony informative sites in each alignment. The number of parsimony informative sites in each alignment was calculated using the “pis” function in PhyKIT. The 2,500 most informative single copy orthologs were concatenated into a single supermatrix using the “create_concat” function in PhyKIT. The resulting alignment had 3,937,461 total sites, 70,307 parsimony informative sites, and 3,799,815 constant sites, which was determined using the “aln_summary” function in BioKIT, v0.0.5 (Steenwyk et al., 2021b), and used as input into IQ-TREE, v2.0.6 (Steenwyk et al., 2021b; Minh et al., 2020). The best fitting substitution model, GTF+F+I+G4 (Tavaré, S., 1986; Vinet &Zhedanov, 2011), was determined using Bayesian Information Criterion in IQ-TREE. Bipartition support was assessed using 5,000 ultrafast bootstrap approximations (Hoang et al., 2018). phylANOVA (Garland et al., 1993; Harmon, et al., 2008; Revell, 2012) test was used to compare gene copy number per orthogroup among environmental and clinical isolates, using a p-value threshold of 0.05. The p-values results are described for each orthogroup at Supplementary Material 2.

## Acknowledgments

We thank Fundação de Amparo à Pesquisa do Estado de São Paulo (FAPESP) 2020/10536-9 (MACH) and 2016/07870-9 (GHG), and Conselho Nacional de Desenvolvimento Científico e Tecnológico (CNPq) 301058/2019-9 and 404735/2018-5 (GHG), both from Brazil, and National Institutes of Health/National Institute of Allergy and Infectious Diseases (R01AI153356), from the USA.

## Figure legends

**Supplementary Figure 1: Comparison of the ontology terms among genes present in CORE and AC orthogroups revealed differences in the abundance of functional classifications between the two main gene groups**. Genes classified by their Gene Ontology terms using PANTHERv16 (Mi et al.,2021). A.Molecular Function; B.Protein Class; C.Hydrolase activity; D.Oxidoreductase activity; right side AC genes, left side CORE genes.

**Supplementary Figure 2: Single nucleotide resolution read depth across A. fumigatus gliotoxin cluster in IFM59779 reveals a deletion overlapping the majority of *glik* cluster**. The Y-axis represents read depth mapped against the Af293 reference gliotoxin cluster (chromosome 6:2346513-2372302). Genes names are labeled and correspond to Afu6g09630 (*gliZ*) and Afu6g09740 (*gliT*). Arrow direction indicates direction of transcription.

**Supplementary Figure 3: Unrooted phylogenetic tree inferred based on single copy genes of all genomes of PanCore analyses**. No significant clade division was observed across the isolates phylogenetic position.

**Supplementary Figure 4: Transposable element *Taf1* and *Aft1* copy number per isolate**. Copy number variation of Aft1 (EF100757.1) and Taf1 (AY971670.1) across isolates (considered nucleotide identity >99.0% to the reference sequence Af293) found Aft1 in 190 isolates Taf1 in 204 isolates varying from 1 to 20 copies; X-axis represents the isolates and the Y-axis the number of sequences identified on genome.

**Supplementary Figure 5: Gene content completeness per isolate**. BUSCO classification presented with assessments for the proportion of genes present in genomes showing the % of complete and single copy genes, % complete and duplicated genes, % of fragmented genes and % of missing genes for all isolates, represented at Y-axis.

**Supplementary Table 1**: **Isolates divergence in terms gene prediction**. Gene number classification of the top 5 biggest and smallest genomes compared to reference isolates Af293 and A1163 (Nierman etal., 2005; Fedorova et al., 2008) showing the variation on gene copy number prediction.

## Notes

### Competing Interest Statement

The authors have declared no competing interest.

