## Supplementary figures and images for "Examination of genome-wide ortholog variation in clinical and environmental isolates of the fungal pathogen *Aspergillus fumigatus*"

### Supplemental Figure 1

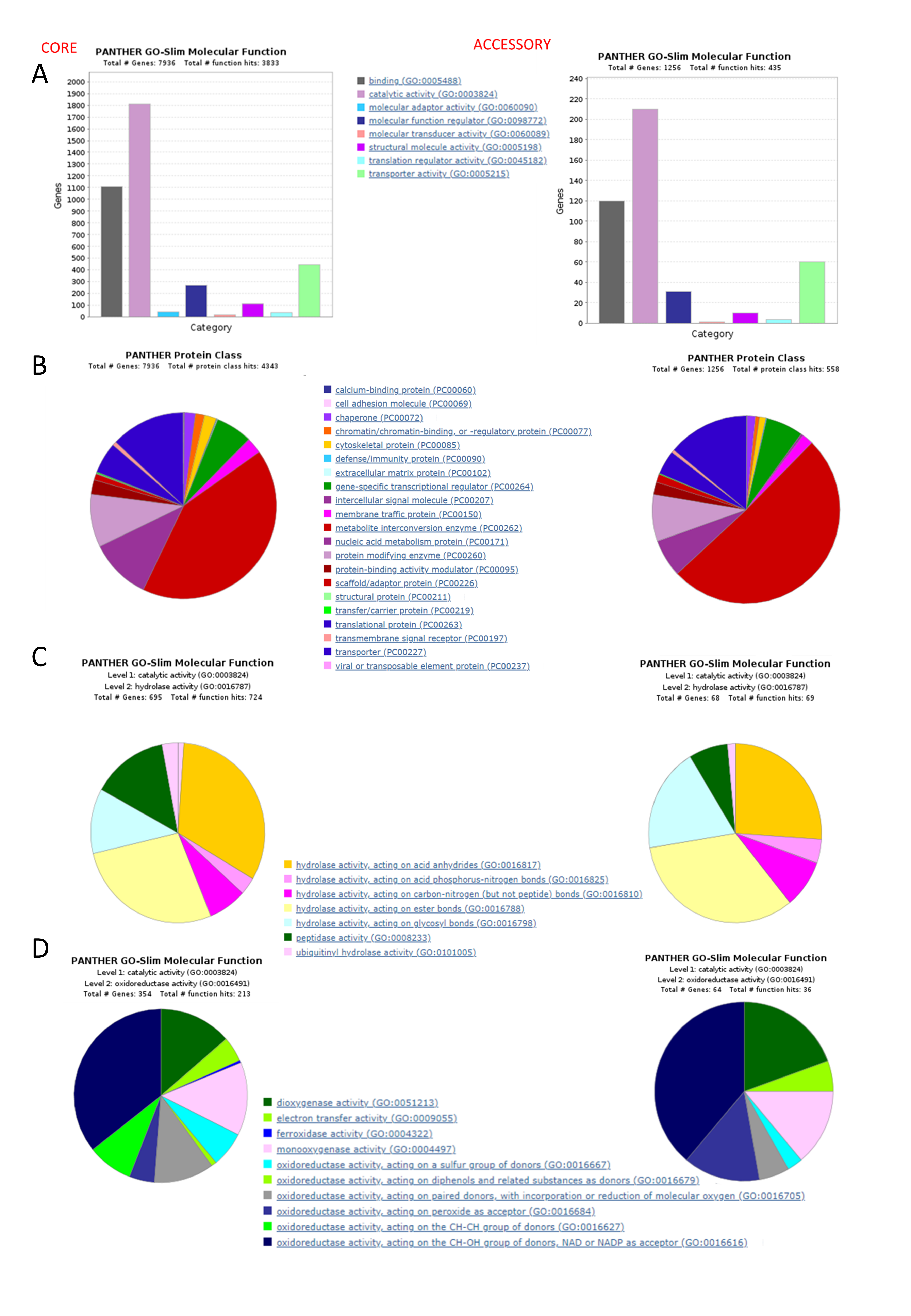

### Supplemental Figure 2

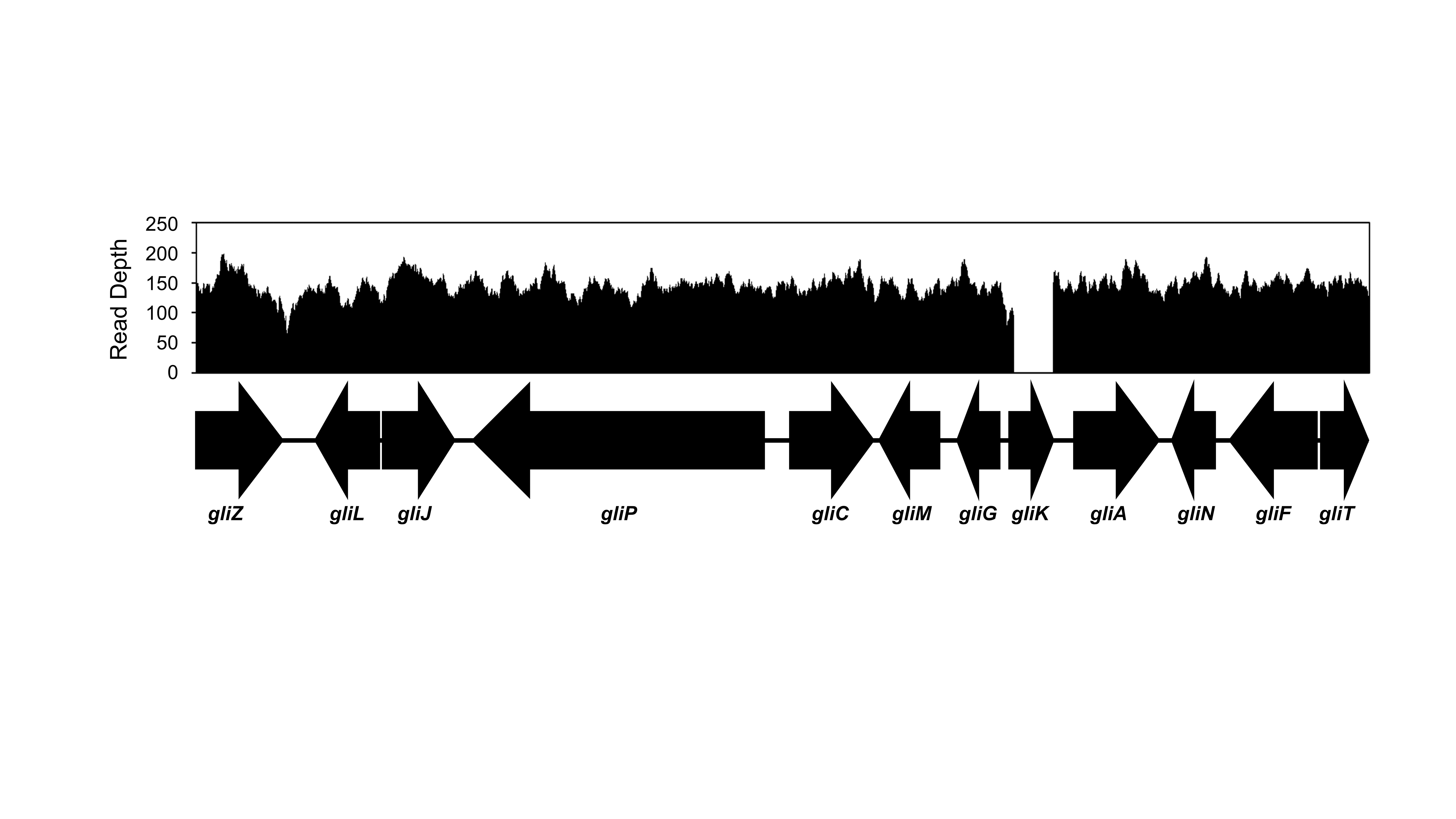

### Supplemental Figure 3

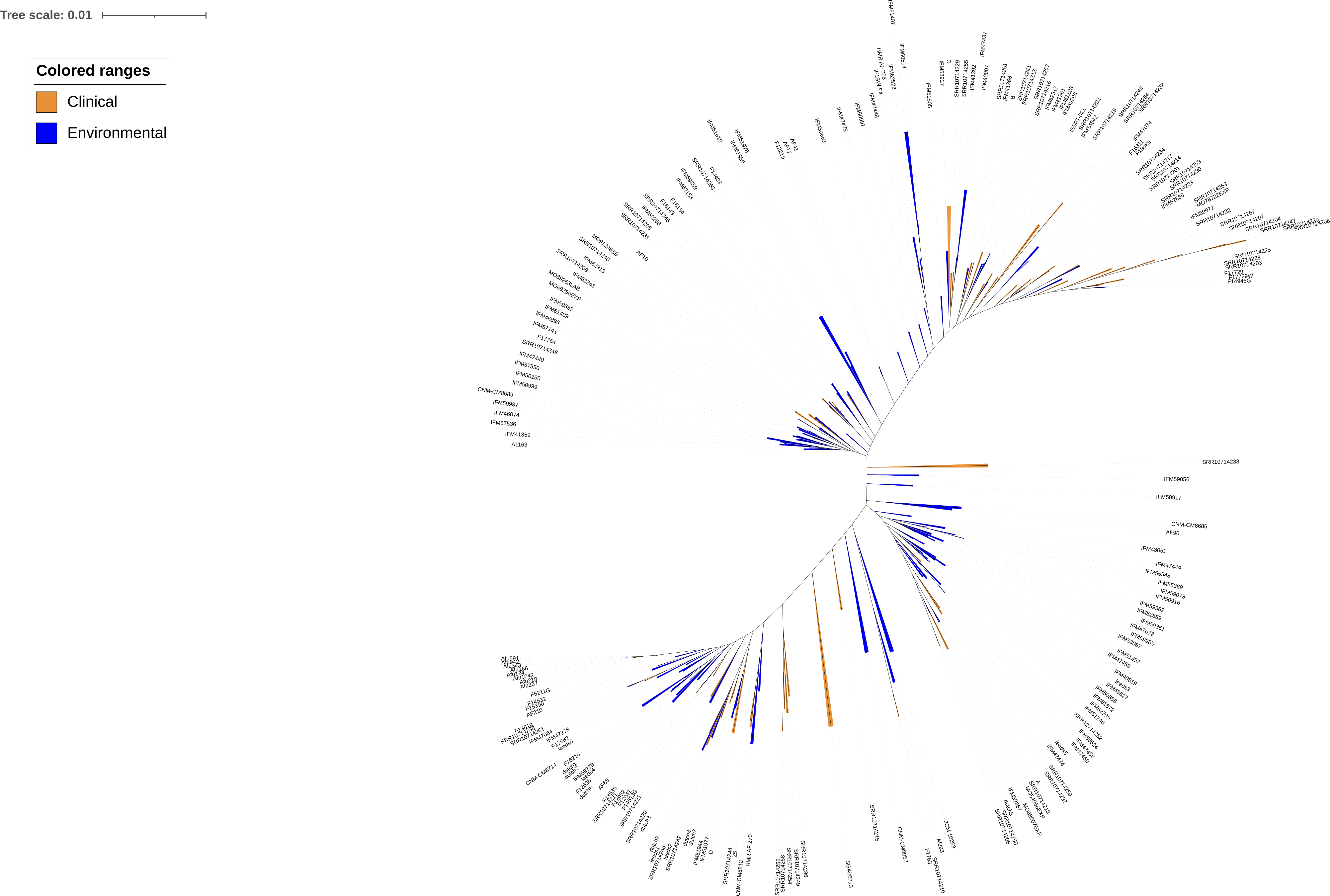

### Supplemental Figure 4

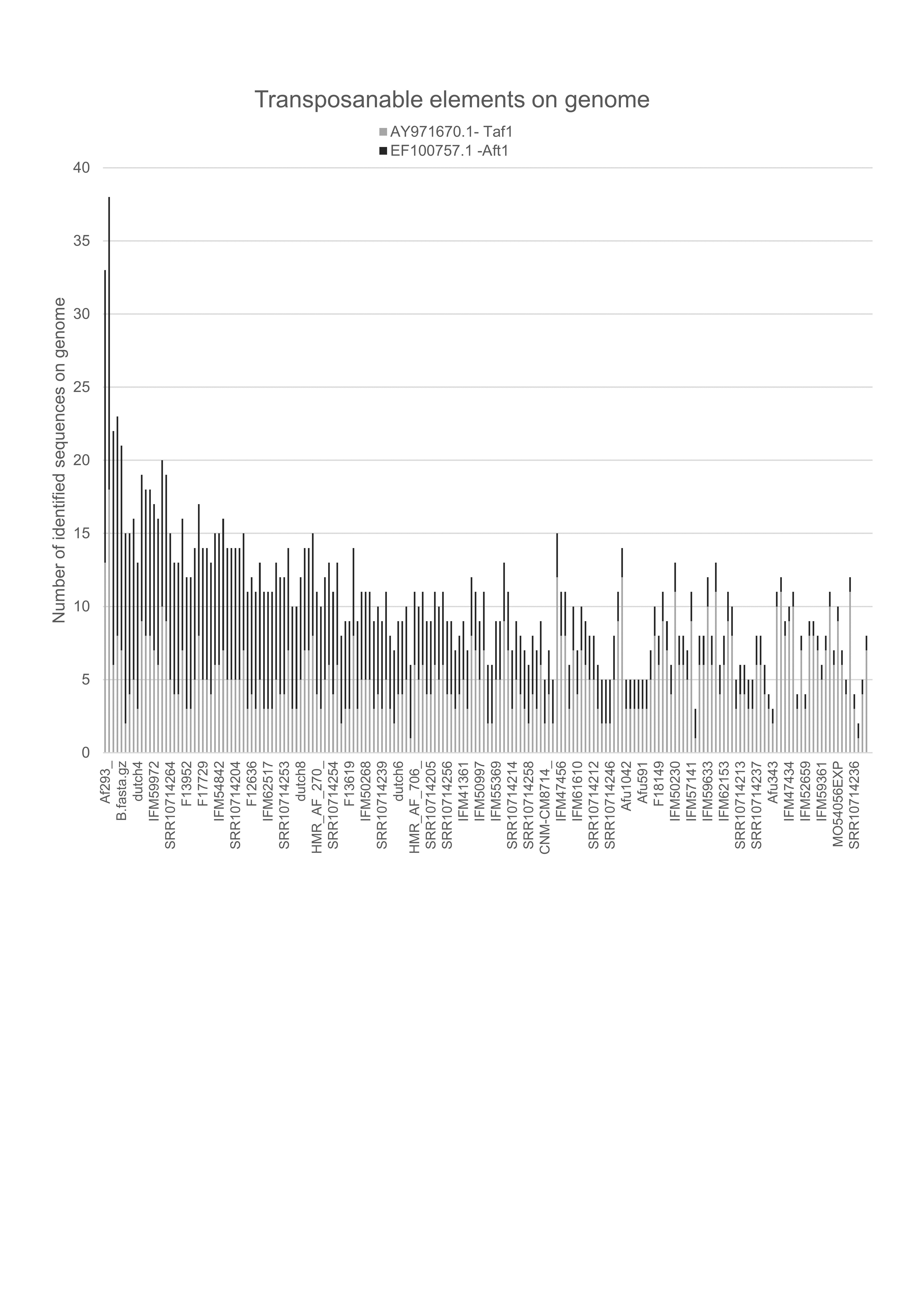

### Supplemental Figure 5

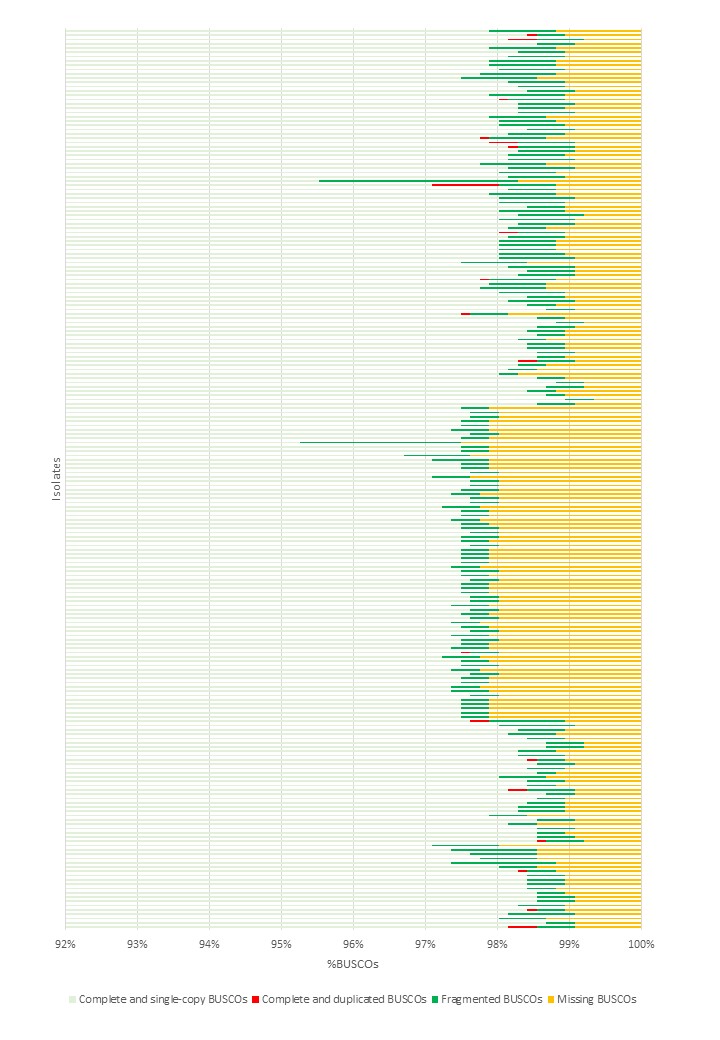
