## Supplemental Table 1 for "Examination of genome-wide ortholog variation in clinical and environmental isolates of the fungal pathogen *Aspergillus fumigatus*"

| Isolate | Predicted sequences on PanCore | Number of Genes on CORE | Number of CORE Orthogroups | Number of genes on ACCESSORY | Number of ACCESSORY Orthogroups | unique genes | Origin |
| --- | --- | --- | --- | --- | --- | --- | --- |
| E-1-65L-4 (SRR10714233) | 9638 | 9144 | 7849 | 7773 | 1295 | 1235 | Environmental |
| B-1-71L-1 (SRR10714232) | 9423 | 9299 | 7830 | 7773 | 1469 | 1411 | Environmental |
| IFM59361 | 9418 | 9385 | 7815 | 7773 | 1570 | 1415 | clinical |
| B-1-26-5 (SRR10714229) | 9413 | 8982 | 7804 | 7773 | 1178 | 1159 | Environmental |
| C-1-72L-1 (SRR10714215) | 9367 | 9288 | 7815 | 7773 | 1473 | 1365 | Environmental |
| A1163_ | 9156 | 9155 | 7794 | 7773 | 1361 | 1262 | Clinical |
| Af293_ | 9094 | 9088 | 7807 | 7773 | 1281 | 1182 | Clinical |
| leeds6 | 8885 | 8885 | 7788 | 7773 | 1097 | 1084 | Clinical |
| IFM48051 | 8883 | 8883 | 7787 | 7773 | 1096 | 1090 | clinical |
| dutch8 | 8879 | 8879 | 7787 | 7773 | 1092 | 1085 | Environmental |
| IFM41392 | 8875 | 8875 | 7786 | 7773 | 1089 | 1084 | clinical |
| leeds4 | 8862 | 8862 | 7784 | 7773 | 1078 | 1073 | Clinical |
| IFM59779 | 8857 | 8856 | 7786 | 7773 | 1070 | 1063 | clinical |

Supplementary Table 1: Gene number classification of the top 5 biggest and smallest genomes compared to reference isolates Af293 and A1163 (Nierman etal., 2005; Fedorova et al., 2008).
